# Associations between meteorological variables, vector indices and dengue hospitalizations in Can Tho, Vietnam: a field survey

**DOI:** 10.1101/664599

**Authors:** Nguyen Phuong Toai, Dang Van Chinh, Nguyen Ngoc Huy, Amy Y. Vittor

## Abstract

**Introduction:** Dengue is a significant cause of morbidity and mortality in Can Tho, a province in the Mekong Delta in Vietnam. In this region, average temperatures have increased by 0.5°C since 1980, and river levels have risen. In a time-series analysis, we previously found that relative humidity was the most important meteorological predictor for dengue hospitalizations in Can Tho. To better understand proximate factors mediating this association, this study examines weather variables in relation to dengue hospitalization rates, vector indices, container productivity and larval elimination and mosquito avoidance behaviors.

**Methods:** Four hundred households were sampled bimonthly for one year in Can Tho. Vector indices of the immature forms of the dengue vector, *Aedes aegypti*, and the productivity of different types of household containers were determined. Dengue hospitalization rates were determined for the study period. Associations between these variables and mean temperature, relative humidity, precipitation, and the number of hours of sun were estimated using mixed effects Poisson regression analysis. Relative productivity of containers was determined by collecting *Ae. aegypti* pupae using a sweep method and adjusting by a calibration factor. *Ae. aegypti* larval density risk factors were determined using multivariate generalized estimating equations with a negative binomial distribution. To examine possible mechanisms mediating the relationship between climate, vectors and dengue, we also interviewed households about mosquito avoidance and larval elimination behaviors.

**Results:** The house-(HI), container-(CI), Breteau (BI), and pupal (PI) indices were associated with relative humidity (1-month lag, IRR_HI_=1.10 (95% CI 1.06, 1.13) per 1% increase), IRR_CI_=1.10 (95% CI 1.02, 1.19), IRR_BI_=1.17 (95% CI 1.14, 1.21), IRR_PI_=1.12 (95% CI 1.10, 1.14)). Vector indices were also associated with precipitation (1-month lag) and to a lesser degree, hours of sun and mean temperature. *Ae. aegypti* larval density was associated with not cleaning water storage containers (RR=2.50, 95% CI 1.59, 3.66), not having access to municipal waste pick-up (RR=3.15, 95% CI2.09, 4.75), disheveled clothes in the home (RR=1.85, 95% CI 1.24, 2.74) and season (RR[rainy season]=3.10, 95% CI 2.18-4.48). The most productive containers were water storage containers (relative pupal productivity 87%). Dengue hospitalization rates were associated with relative humidity (2-month lag, IRR=1.11 (95% CI 1.06, 1.17) per 1% increase). Only the PI (1-month lag) was significantly associated with dengue hospitalization rates (IRR 1.04, 95% CI 1.00, 1.07). Mosquito avoidance behaviors were more frequent in the dry season (92.5% vs. 86.0% of interviewees endorsed one or more forms of mosquito prevention, p<0.001). There was also less use of larval elimination strategies (39.2% vs. 50.5%, p<0.001) during the rainy versus the dry season.

**Conclusion:** Our study reveals a strong effect of relative humidity on vector indices and dengue hospitalization rates. This may be due to the mosquito’s vulnerability to desiccation, and the association warrants further study. Our findings also demonstrate, however, that during the rainy season when mosquito prevention is most needed, the use of fans, repellant coils and maintenance of water storage containers is actually reduced. Water storage containers were by far the most productive of pupae, and should be targeted in vector control activities.

**Author summary:** Climate plays an important role in the geographic distribution and burden of disease due to dengue, owing to the vector and virus’ sensitivity to temperature, humidity, and rainfall. In the Mekong Delta in Vietnam, where dengue poses a significant health burden, average temperatures have increased by 0.5°C since 1980. To better understand the influence of climate on dengue, this study examines its influence on dengue hospitalization rates, vector breeding behavior and human mosquito avoidance behaviors. We sampled 400 households every 2 months for one year for the presence of the dengue vector, *Aedes aegypti*, and the productivity of different types of household containers. Human mosquito avoidance behaviors, such as the use of fans, mosquito repellant, and larval elimination strategies were also recorded. The association between dengue hospitalizations, mean temperature, relative humidity, precipitation, and the number of hours of sun were established, and risk factors for the abundance of *Ae. aegypti* larvae were determined. We found that relative humidity is positively associated with the presence of *Ae. aegypti* immature forms, and that large jars used for water storage serve as the most important source of this vector. We also determined that people engage in mosquito avoidance/larval elimination strategies more frequently in the dry season versus the rainy season, despite increased vector breeding and dengue hospitalizations during the rainy season. This temporal disconnect between peak vector activity and dengue hospitalization rates vis-à-vis mosquito control strategies is a potential area for intervention.

## Introduction

Dengue fever is caused by one of four serotypes of dengue virus (family *Flaviviridae*, genus *flavivirus*), a single-stranded positive-sense RNA virus. It is transmitted by *Aedes* species mosquitoes and usually causes a self-limited febrile illness (classic dengue fever), characterized by fever, headache, retro-orbital pain, arthralgia, myalgia, and rash. Severe forms of dengue (dengue hemorrhagic fever and dengue shock syndrome) are rare, but disproportionately affect young children and may result in death. In the past several decades, the geographic range of dengue has expanded greatly and dengue is now endemic throughout the subtropics and tropics. Current estimates place yearly incidence at approximately 390 million cases[1]. The underlying causes for this expansion are thought to be due to increased human mobility, poorly planned urbanization, the breakdown of vector control programs, the lack of public health infrastructure, and climate change [2, 3].

In Vietnam, approximately 125,000 dengue cases occur yearly[4], and this disease accounts for a large portion of hospitalizations[4]. Over 70% of cases occur in the southern region of the country [4]. The city-province of Can Tho lies in this southern region on the Mekong Delta, and is subject to frequent flooding. Climate projections have estimated that by 2030, the business district of the city will be submerged under 50cm of water during the peak rainy season[5]. Furthermore, the average air temperature in the region has increased by 0.5°C since 1980, with a projected increase of 1.1-1.4°C by 2050[5]. There is strong motivation on part of the city leadership to understand the health effects of such projections. Since climate plays an important role in the geographic and temporal distribution of dengue[2, 6], it is important to gain a better understanding of the ways in which the vector responds to climate. In addition, understanding associations between behavioral elements (e.g. water storage habits, use of mosquito avoidance measures) and climate will provide important insights into the human landscape and possible intervention strategies.

Previously, we reported on the associations between dengue hospitalizations in this region and climate between 2004 and 2011 in a time-series analysis[7]. We found that the dengue hospitalization rate in Can Tho was significantly associated with relative humidity with a lag of one month. To better understand the entomological and behavioral factors that may be contributing to this association, we conducted this prospective study. Specifically, we analyze indices of immature forms of *Ae. aegypti*, dengue hospitalization rates, container productivity, and mosquito avoidance/larval elimination behaviors by weather variables.

## METHODS

### Study setting and design

The study was conducted in two districts in Can Tho from June 2012 to June 2013 (Figure 1). The City of Can Tho (10.0333°N, 105.7833°E) lies on the Hau River in the Mekong Delta and is a regional hub for commerce, education, and culture. It has a population of 1.2 million with a density of 868 people/km^2^[8]. The elevation throughout the city ranges from 0.8 to 1.5m above sea level. Can Tho’s climate is tropical and monsoonal with an average annual temperature of 27°C. The rainy season usually occurs from May to November and provides 90% of the yearly rainfall, which averages 1600-2000mm. There are nine districts in Can Tho, which range from urban to rural. However, urbanization has been occurring at a rapid pace. Between 1999 and 2009, the urban population grew by 41.5%[9]. The two districts selected for this study are Ninh Kieu and Binh Thuy (Fig 1). The former lies in the heart of Can Tho city and is urban, while the latter is suburban/rural.

**Fig 1.**
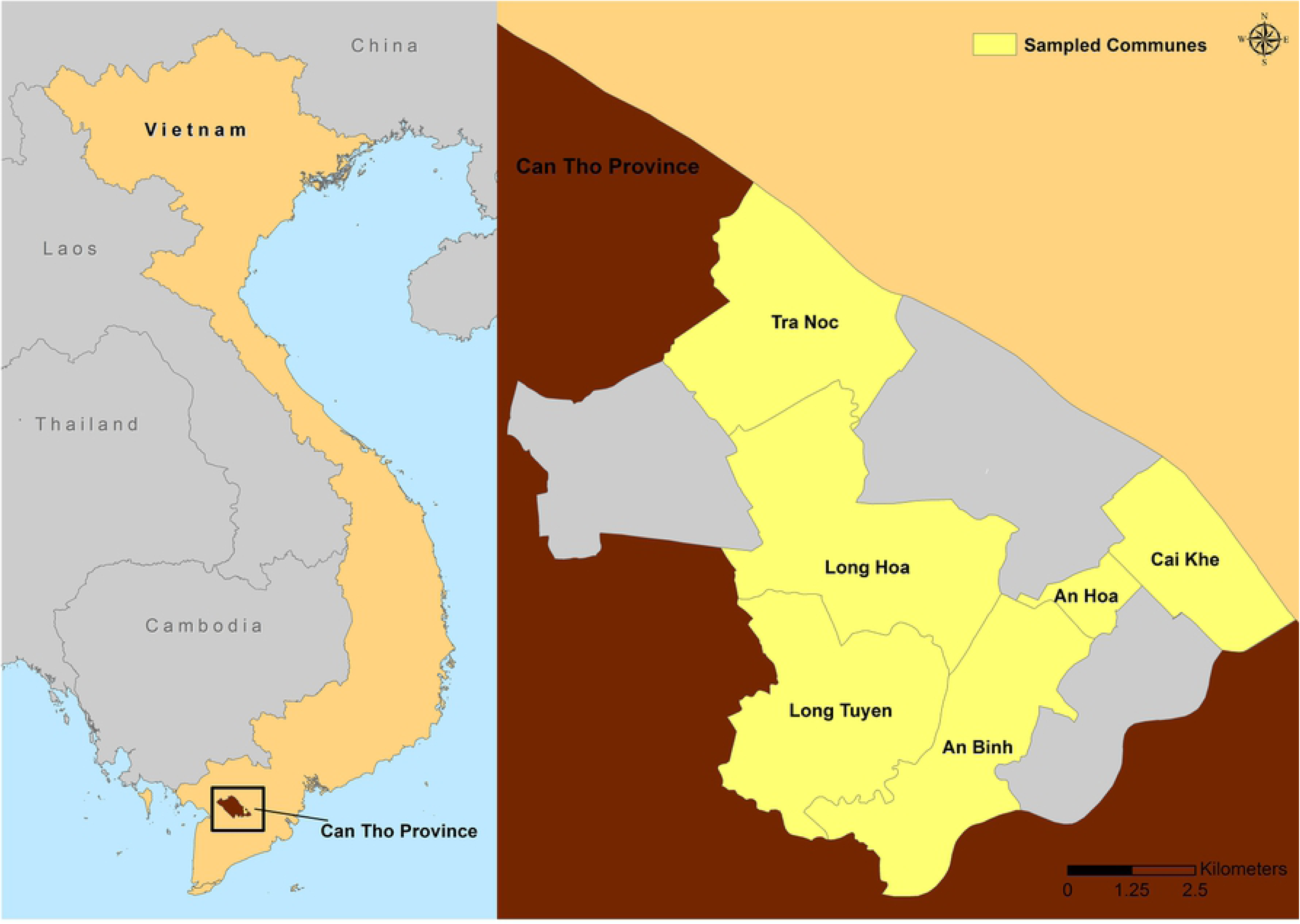
Map of the study area, illustrating the six wards in the province of Can Tho that were sampled between June 2012 and June 2013. Base layers were provided by the Department of Information and Technology of Can Tho City in Can Tho City (http://stnmt.cantho.gov.vn/ttqt). Cai Khe, An Hoa, and An Binh (control) wards are in Ninh Kieu district; Long Hoa, Long Tuyen, and Tra Noc (control) wards are in Binh Thuy district.

Two wards in each of the two districts were selected for the study (Ninh Kieu district: Cai Khe (popn 33,018) and An Hoa (popn 22,367) wards; Binh Thuy district: Long Hoa (popn 13,471) and Long Tuyen (popn 13,250) wards). A third ward in each district was sampled at the beginning and the end of the study as controls for the effect of visitation of study staff on larval abundance (Ninh Kieu district: An Binh ward (popn 30,041); Binh Thuy district: Tra Noc ward (popn 10,513)). One hundred households were randomly sampled within each study sector. 93.5 % of households approached agreed to participate in the study. Once consented, households were visited once every two months. At each visit, larval and pupal collections were conducted in conjunction with a survey of demographics, mosquito avoidance behaviors, and larval elimination strategies.

### Data collection

#### Entomological survey

Larval and pupal surveys were conducted by fieldworkers sampling all water-containing elements in the selected households and the peridomestic environment. Container type and quantity were noted, and mosquito larvae and pupae were quantified. Larvae and pupae in small containers (<20L) were removed using a pipette, while immature mosquito stages were sampled in large containers (>20L) by using sweep nets according to the pupal survey method described in[10, 11]. Larvae were identified by entomological personnel at the Can Tho Preventive Medicine Center, while pupae were allowed to emerge and were then identified as adult mosquitoes. Various entomological indices were derived. The Breteau index (BI) is the number of positive containers per 100 houses sampled. The house index (HI) is the percentage of houses infested with larvae or pupae. The container index (CI) is the percentage of water-holding containers infested with larvae or pupae. The pupal index (PI) is the number of pupae per 100 households. The larval density (LD) was defined as the number of larvae per household. To assess the productivity of different container types, relative pupal productivity was calculated by dividing the number of pupae in a given container by the total number of pupae in all containers in the study area as described in [12].

#### Dengue hospitalizations

Monthly counts of dengue hospitalizations by study ward were obtained from the Can Tho Preventive Medicine Center. Case reporting to the health department is compulsory. Private clinics diagnosing severe dengue are required to send patients to government-run district- and province hospitals. Dengue is diagnosed by NS1 antigen tests in Can Tho’s hospitals, and 7% of NS1 antigen positive cases are further tested by MAC ELISA, and 3% are tested by RT-PCR. Monthly hospitalization rates were determined by using the 2009 populations for each ward.

#### Household survey

The survey was administered to consenting residents, and featured questions pertaining to demographics, socioeconomics, health, and water storage and mosquito avoidance behavior. The survey was administered to each household bimonthly for the duration of the study.

#### Meteorological data

Monthly mean temperatures, relative humidity, precipitation data, and hours of sunshine were obtained from the Can Tho Meteorological Station.

### Statistical analysis

Data was entered into EpiData (Lauritsen 2008), and analyzed using STATA v14 (College Station, TX, USA). To explore the seasonal differences in vector indices and dengue rates, two-sample t-tests were first performed. Subsequently, the effects of individual meteorological factors on vector indices and dengue rates were examined using a mixed-effects Poisson regression. A mixed effects model was employed to account for ward-level variability (random effects) when estimating the association of meteorological variables (fixed effects). The possibility of a time lag (0-, 1-, 2-months) was explored by separately modeling each lag. The meteorological variables were highly correlated. To avoid issues related to collinearity, separate models were constructed for each meteorological variable and each outcome variable (vector indices and dengue hospitalization rates). The models were further examined by comparing random-slope and random intercept models using likelihood ratio tests. Akaike’s information criterion (AIC) was employed for model selection.

Risk factors for *Ae. aegypti* larval density were modeled using generalized estimating equations (GEE) with a negative binomial distribution. Variables with a p-value of less than 0.25 in the univariate analysis were included in the model in a stepwise fashion, and maintained in the model if significant at the 0.05 level.

Larval and pupal indices and counts were used to determine container productivity. Since jars and cement tanks hold large volumes of water, the absolute count of larvae and pupae collected in a sweep in these containers was adjusted by a calibration factor (C-2Fs) assuming that water level is at two thirds the total capacity, as described in [13]. Mosquito avoidance and larval elimination behaviors were analyzed by season using a McNemar test for paired proportions. The maps were created using ArcMap 10.2.2 software by ESRI, using base layers provided by the Department of Information and Technology of Can Tho City in Can Tho City (http://stnmt.cantho.gov.vn/ttqt).

### Ethical considerations

The study was approved by the Can Tho Department of Health Committee for Human Research (COA. No. 246/SYT, protocol No. 052/NCKH-SYT). Study participation was voluntary, and all adult participants provided informed consent. No children participated in this study.

## RESULTS

### Study participant and meteorological characteristics

The households (n=400) were surveyed 7 times each over the course of 13 months. Additional control households (n=200) were sampled the first and last month of the study. Women comprised 63.5% of those interviewed, and most participants were over 45 years of age (Table 1). The majority of participants had completed secondary education or higher (64.3%), and literacy levels were high. The availability of tap water and municipal waste pickups were good (88.3% and 70.3%, respectively), but not universal. It is also important to note that while tap water was widely available, frequent water shortages meant that tap water could not be relied upon continuously. Most households had between 4 and 6 inhabitants.

**Table 1.** Study respondent characteristics (n=400 households).

| Characteristics |  | Frequency (n) | % |
| --- | --- | --- | --- |
| Age | 15-24 yrs | 29 | 7.3 |
|  | 25-34 yrs | 45 | 11.3 |
|  | 35-44 yrs | 79 | 19.8 |
|  | > 45 yrs | 247 | 61.8 |
| Sex | Male | 146 | 36.5 |
|  | Female | 121 | 63.5 |
| Literacy | Illiterate | 17 | 4.3 |
|  | Literate | 383 | 95.8 |
| Education | Less than high school | 143 | 35.8 |
|  | High school graduate | 201 | 50.3 |
|  | College graduate | 56 | 14.0 |
| Piped water supply | No | 46 | 11.5 |
|  | Yes | 353 | 88.3 |
| Municipal waste | No | 118 | 29.5 |
|  | Yes | 281 | 70.3 |
| No. inhabitants in the home | 1-3 | 111 | 27.8 |
|  | 4-6 | 236 | 59.0 |
|  | >=7 | 53 | 13.3 |

Mean monthly temperatures were very constant throughout the study period, varying only by 1°C around the mean temperature of 27.7°C (Figure 2). Cumulative annual rainfall was 1135mm. Most of the precipitation occurred between May and October, accounting for 91% of the yearly total. Relative humidity was high throughout the year (annual mean 82%), ranging between 77% during the dry season and 88% during the rainy season. The monthly hours of sunshine ranged between 210 to 294 hours during the dry season, and 148 to 251 hours in the rainy season.

**Fig 2.**
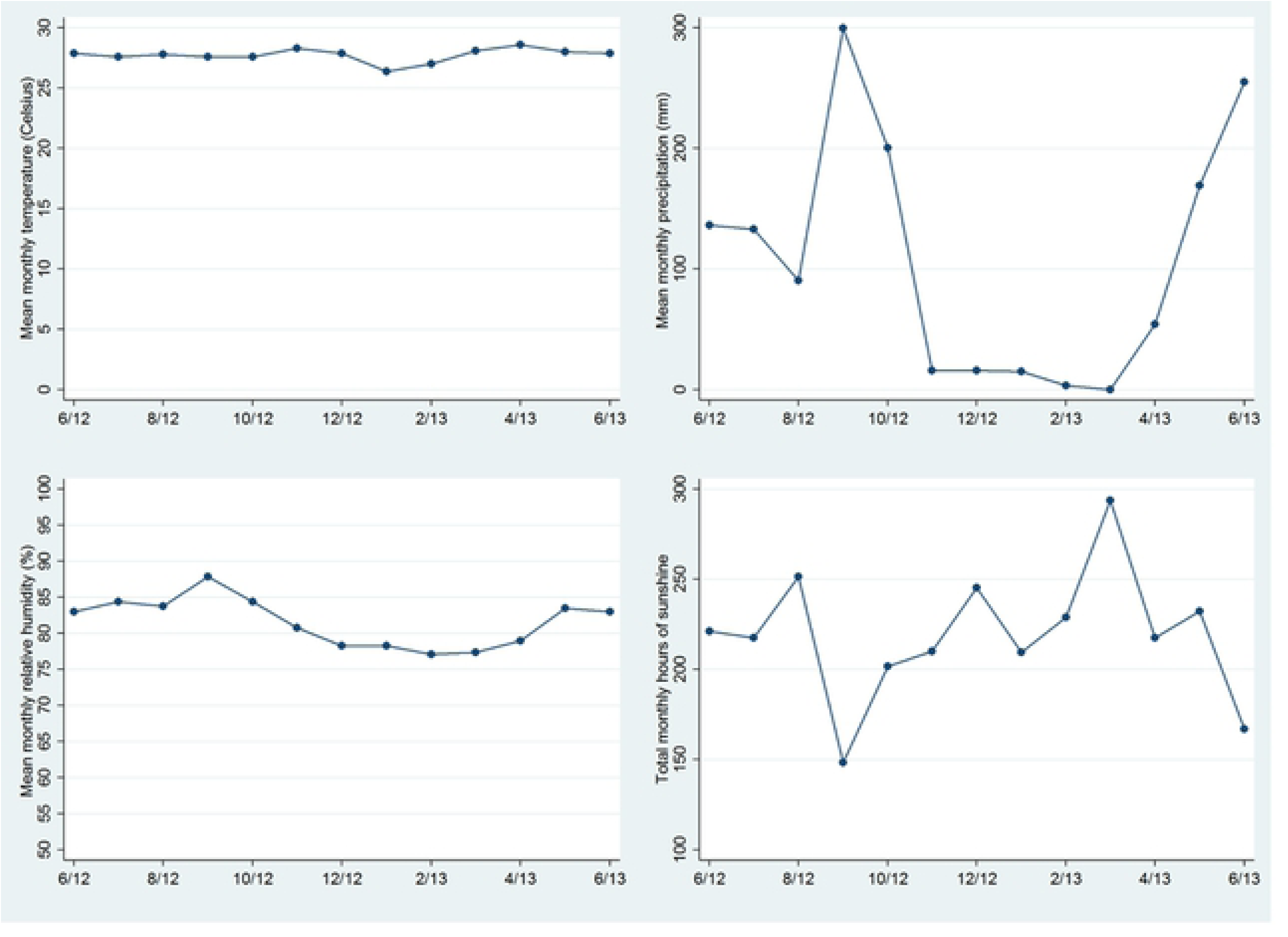
Mean monthly temperature, precipitation, relative humidity, and hours of sun in Can Tho, Vietnam.

#### *Aedes aegypti* vector indices

All vector indices were markedly higher during the rainy season (Figure 3). The house index (HI) ranged from 7.5% (95% CI 4.0, 11.0) in the dry season to 14.2% (10.5, 17.8) in the rainy season (p=0.01). The container index (CI) was 1.3% (95% CI 0.8, 1.9) in the dry season and 2.8% (95% CI 2.0, 3.6) in the rainy season (p=0.01). The Breteau index (BI) was 8.3 (95% CI 4.0, 12.7) in the dry season, and 21.7 (95% CI 12.9, 30.5) in the rainy season (p=0.03), and the pupal index (PI) was (95% CI 11.9, 32.0) in the dry season and 72.8 (95% CI 36.6, 108.9) in the rainy season (p=0.03). There was a single outlier for PI, however. All PI values were between 0 and 111 with the exception of a single observation at 368. The mean larval density (number of larvae per household) (LD) was 2.2 (95% CI 1.7, 2.8) in the dry season and 6.7 (95% CI 5.5, 7.8) in the rainy season (p<0.001). HI and CI were closely correlated (ρ=0.81), and these were in turn moderately to strongly correlated with the Breteau index (ρ_HI_=0.82, ρ_CI_=0.69).

**Fig 3.**
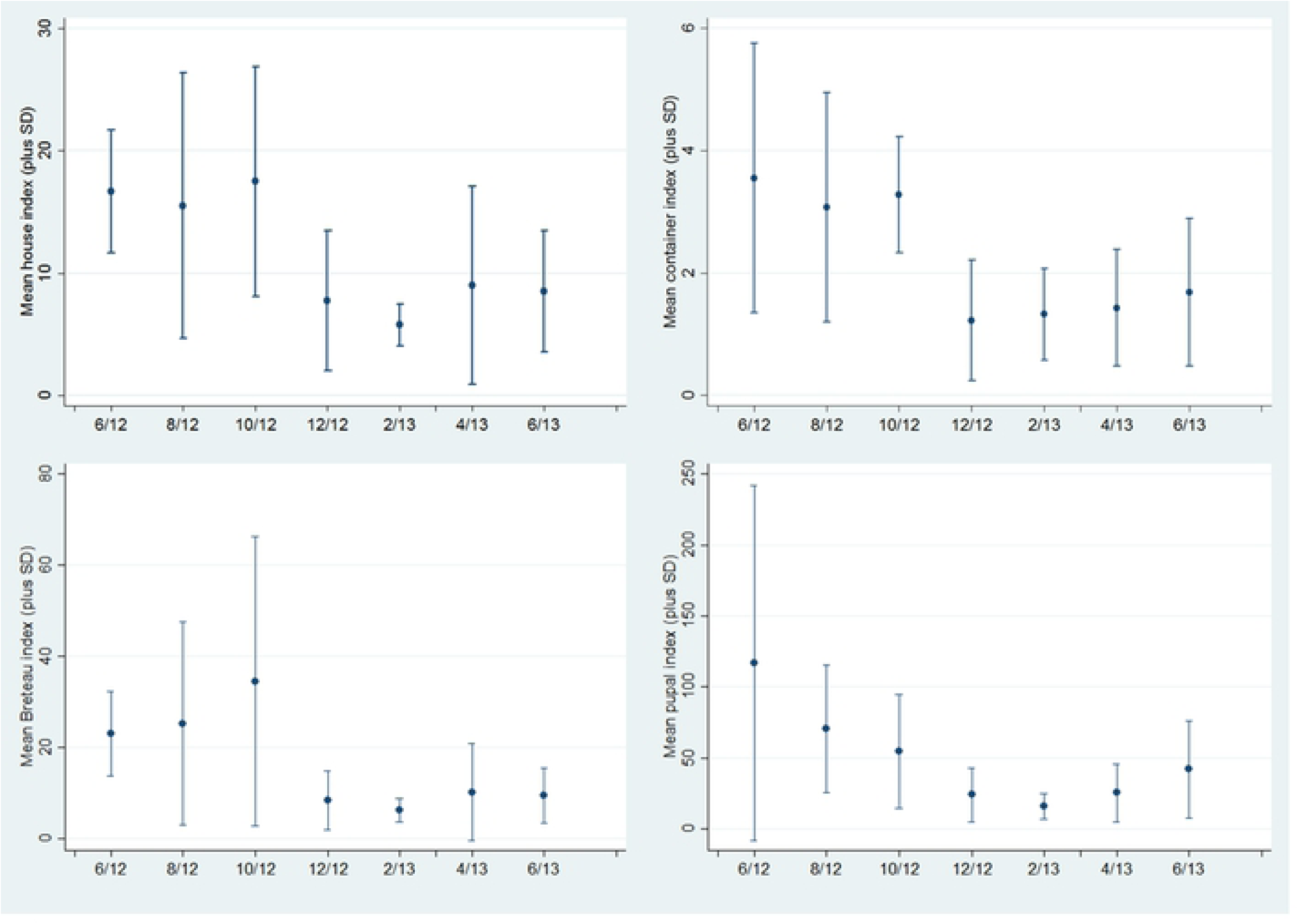
Vector indices (house index, container index, Breteau index, pupal index) during the study period in Can Tho, Vietnam.

To examine the association between vector indices and meteorological factors in more detail, mixed effects Poisson regression coefficients were estimated for 0-, 1-, and 2-month lags (Table 2). The most pronounced associations were seen with relative humidity. Relative humidity was positively associated with virtually all vector indices at all lags. Precipitation was also strongly associated with all indices at 1-month lag, and variably associated at 0- and 2-month lags. The monthly hours of sunshine were negative associated with HI, BI, and PI at 1-month lag. Temperature was significantly correlated only with PI, and most strongly at a lag of 2-months. PI was the only vector index that demonstrated a significant association with all four meteorological factors. AIC were generated for each model. The 1-month lag models had the lowest AIC and thus the best fit compared to the 0- and 2-month lag models.

**Table 2.** Univariate mixed effects Poisson regression incidence rate ratios for vector indices and meteorological factors with 0-, 1- and 2-month lags. Bolded entries are statistically significant.

|  | Lag (months) | House index (IRR, 95% CI) | Container index (IRR, 95% CI) | Breteau index (IRR, 95% CI) | Pupal index (IRR, 95% CI) |
| --- | --- | --- | --- | --- | --- |
| Mean monthly temp (°C) | 0 | 1.07<br>(0.84, 1.38) | 0.95<br>(0.54, 1.67) | 0.93<br>(0.75, 1.14) | **1.18\***<br>**(1.05, 1.32)** |
|  | 1 | 1.07<br>(0.87, 1.31) | 0.96<br>(0.61, 1.50) | 1.02<br>(0.86, 1.20) | **1.14\***<br>**(1.02, 1.27)** |
|  | 2 | 0.96<br>(0.74, 1.24) | 1.09<br>(0.63, 1.89) | 0.98<br>(0.79, 1.21) | **1.33\*\***<br>**(1.16, 1.52)** |
| Precipitation (cm/month) | 0 | **1.01\***<br>**(1.00, 1.03)** | 1.02<br>(0.99, 1.04) | **1.02\*\***<br>**(1.01, 1.03)** | **1.02\*\***<br>**(1.02, 1.03)** |
|  | 1 | **1.03\*\***<br>**(1.02, 1.04)** | **1.03\***<br>**(1.00, 1.06)** | **1.05\*\***<br>**(1.04, 1.06)** | **1.03\*\***<br>**(1.02, 1.04)** |
|  | 2 | 1.02<br>(1.00, 1.04) | 1.01<br>(0.97, 1.06) | **1.02\*\***<br>**(1.01, 1.04)** | **1.02\*\***<br>**(1.01, 1.03)** |
| Mean monthly | 0 | **1.14\*\*** | **1.15\*** | **1.23\*\*** | **1.22\*\*** |
| relative humidity (%) |  | (1.09, 1.19) | (1.04, 1.28) | (1.18, 1.28) | (1.19, 1.25) |
|  | 1 | 1.10**<br>(1.06, 1.13) | 1.10*<br>(1.02, 1.19) | 1.17**<br>(1.14, 1.21) | 1.12**<br>(1.10, 1.14) |
|  | 2 | 1.09**<br>(1.04, 1.14) | 1.08<br>(0.98, 1.19) | 1.16**<br>(1.12, 1.21) | 1.10**<br>(1.07, 1.12) |
| Sunshine (10hrs/month increments) | 0 | 1.02<br>(0.99, 1.07) | 1.02<br>(0.94, 1.11) | 1.04*<br>(1.00, 1.07) | 1.02*<br>(1.01, 1.04) |
|  | 1 | 0.95**<br>(0.92, 0.98) | 0.94<br>(0.88, 1.01) | 0.90**<br>(0.88, 0.92) | 0.96**<br>(0.94, 0.97) |
|  | 2 | 1.09*<br>(1.01, 1.17) | 1.11<br>(0.94, 1.31) | 1.20**<br>(1.13, 1.28) | 1.02<br>(0.98, 1.06) |

\* p<0.05
\*\* p≤0.001

The two communes that were only sampled the first and last months of the study period (June 2012 and June 2013) demonstrated a lesser rate of decline in vector indices between these two time points compared to the regularly sampled communes. The ratio of HI (first month/last month) for the two communes sampled twice was 2.1 compared to 2.8 (p=0.36) for the regularly sampled communes. Similarly, the CI (first month/last month) was 2.0 vs 2.4 (p=0.37), the BI (first month/last month) was 2.8 vs 2.9 (p=0.48), the PI 1.5 versus 19.8 (p=0.27), and the LD (first month/last month) was 2.2 vs 3.9 (p=0.22), for the twice-sampled communes compared to the regularly sampled communes, respectively. While these results are not statistically significant, the effect of sampling monthly or bimonthly may have impacted the outcome by reducing the abundance of *Ae. aegypti.*

### Risk factors for *Ae. aegypti* larval density

Multivariate analysis of risk factors associated with *Ae. aegypti* larval density demonstrated that larval density was positively associated with the rainy season (RR 3.1, 95% CI 2.18 – 4.48), not cleaning water containers (RR 2.5, 95% CI 1.59 – 3.66), not having municipal waste management (RR 3.15, 95% CI 2.09 – 4.75), and having clothes disorganized in the home (RR 1.85, 95% CI 1.24 – 2.74) (Table 3). Other behaviors and weather variables were not significantly associated with larval abundance upon controlling for these four factors.

**Table 3.**
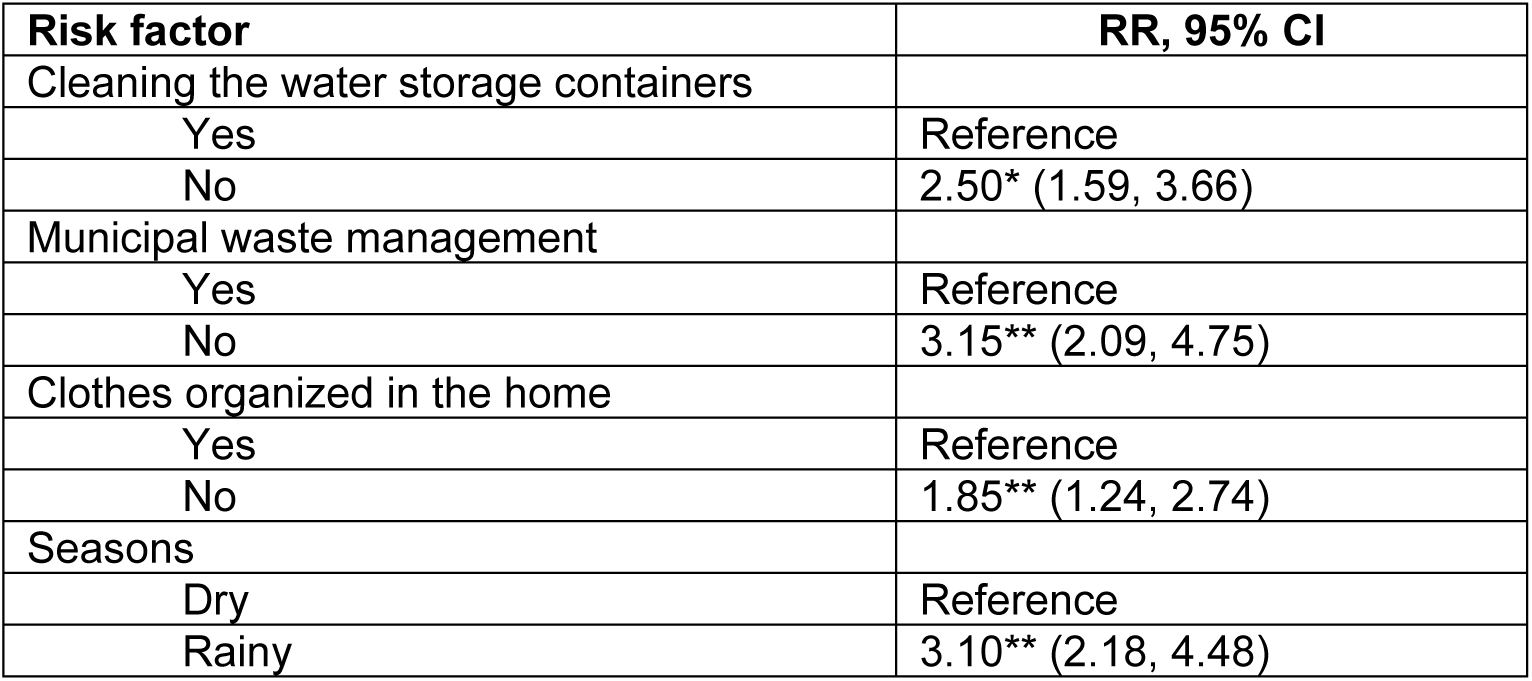
Multivariate model of risk factors for *Aedes aegypti* larval abundance using generalized estimating equations.

### Relative productivity of containers

Indoor and outdoor household containers were sampled for *Ae. aegypti* larvae and pupae. In the rainy season, 14,240 containers were sampled (Table 4a). In the dry season, 6,291 containers were sampled (Table 4b). A total of 1,688 *Ae.aegypti* pupae were collected in the rainy season, with dramatically less in the dry season (263 pupae). Since water storage containers and cement tanks hold large volumes of water, the absolute count of pupae collected in a sweep was adjusted by a calibration factor (C-2Fs) assuming that water level is at two-thirds the total capacity[13]. In both seasons, the most productive containers were the water storage containers with a relative productivity of 87% and 88% in the rainy and dry seasons, respectively. This was followed by cement tanks (relative productivity 10% and 7% in the rainy and dry seasons). Other containers such as buckets, vases, and miscellaneous other containers were common, but none of these yielded many larvae nor pupae relative to the jars.

**Table 4.**
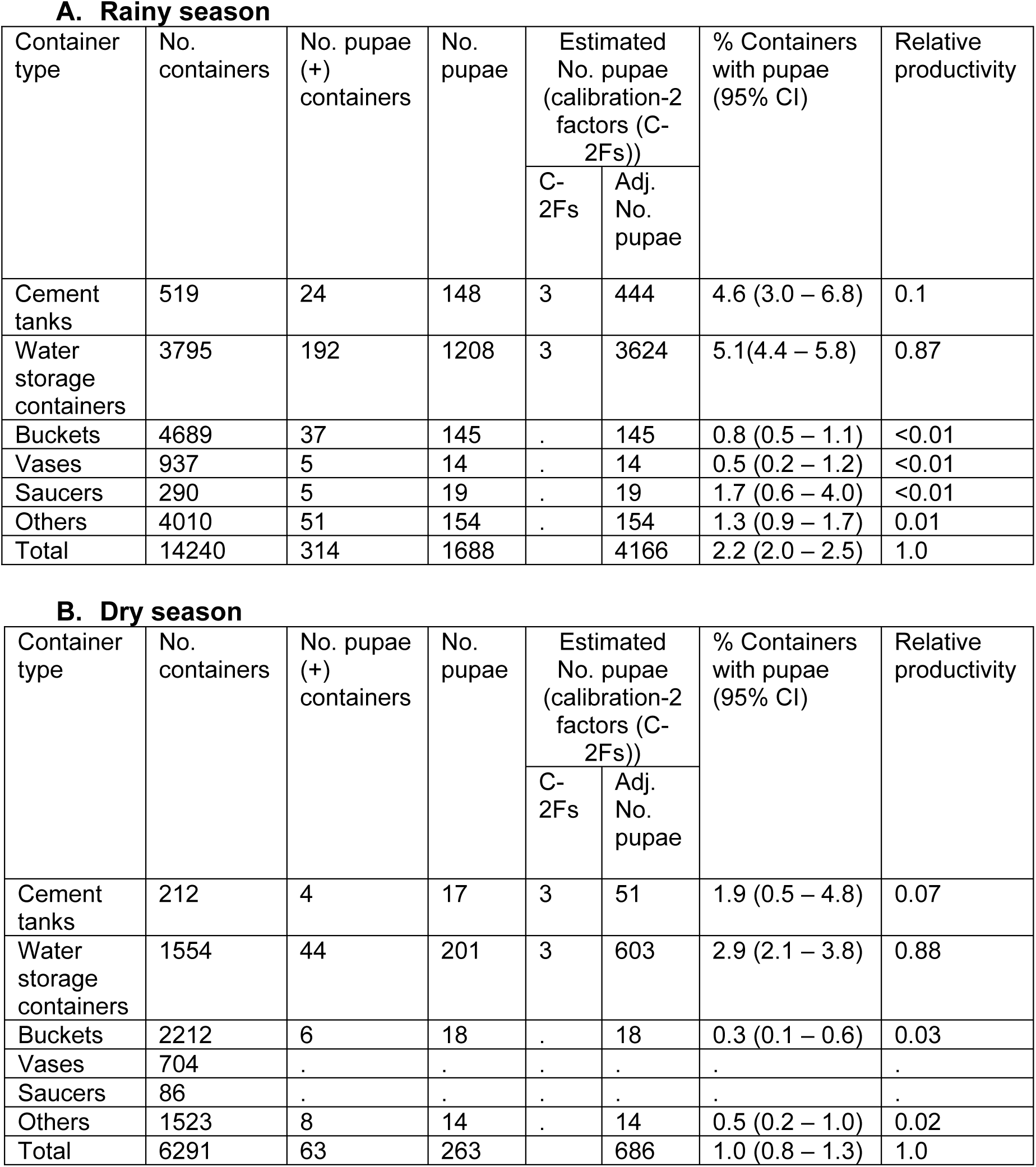
A) Container pupal productivity in the rainy season, and B) container pupal productivity in the dry season.

### Dengue hospitalization rates

The dengue hospitalization rate varied by ward and by month (Figure 4). Long Tuyen and An Hoa wards had the highest dengue hospitalization rates, averaging 15.7 (95%CI 9.9, 21.4) and 15.5 (95%CI 7.6, 23.3) hospitalizations/month/100,000 population, respectively (Table 5). The other wards had rates similar to one another, averaging 5.9 to 13.2 hospitalizations/month/100,000 population. Peak hospitalizations occurred in August and December 2012 (17.9 and 18.8 hospitalizations/month/100,000 population, respectively) and nadirs occurred in February and May 2013 (2.4 and 3.3 hospitalizations/month/100,000 population, respectively).

**Table 5.** Univariate mixed effects Poisson regression incidence rate ratios for dengue hospitalization rates, A) meteorological factors, and B) vector indices with 0-, 1- and 2-month lags.

| **A** | Lag | Mean monthly temperature (°C) | Precipitation (cm/month) | Mean monthly relative humidity (%) | Sunshine (10hr/month increments) |
| --- | --- | --- | --- | --- | --- |
| Dengue monthly hospitalization rate (per 100,000) | 0 | 1.31<br>(0.96, 1.80) | 1.01<br>(0.99, 1.03) | **1.07\***<br>(**1.02, 1.13**) | 0.98<br>(0.93, 1.02) |
|  | 1 | **1.38\***<br>(**1.00, 1.91**) | 1.01<br>(1.00, 1.03) | **1.10\*\***<br>(**1.04, 1.15**) | 0.99<br>(0.94, 1.04) |
|  | 2 | 0.98<br>(0.73, 1.33) | **1.04\*\***<br>(**1.02, 1.05**) | **1.11\*\***<br>(**1.06, 1.17**) | **0.94\***<br>(**0.90, 0.99**) |

| **B** | Lag | HI | CI | BI (10 unit increments) | PI 10 unit increments) |
| --- | --- | --- | --- | --- | --- |
| Dengue monthly hospitalization | 0 | 1.01<br>(0.95, 1.07) | 1.04<br>(0.80, 1.36) | 1.07<br>(0.95, 1.20) | 1.02<br>(0.91, 1.13) |
|  | 1 | 1.03 | 1.09 | 1.14 | **1.04\*** |
| rate (per 100,000) |  | (0.99, 1.07) | (0.92, 1.30) | (0.99, 1.30) | **(1.00, 1.07)** |
|  | 2 | 1.01<br>(0.98, 1.05) | 1.02<br>(0.87, 1.18) | 1.10<br>(0.97, 1.25) | 1.01<br>(0.98, 1.04) |

\* p<0.05
\*\* p≤0.001

**Fig 4.**
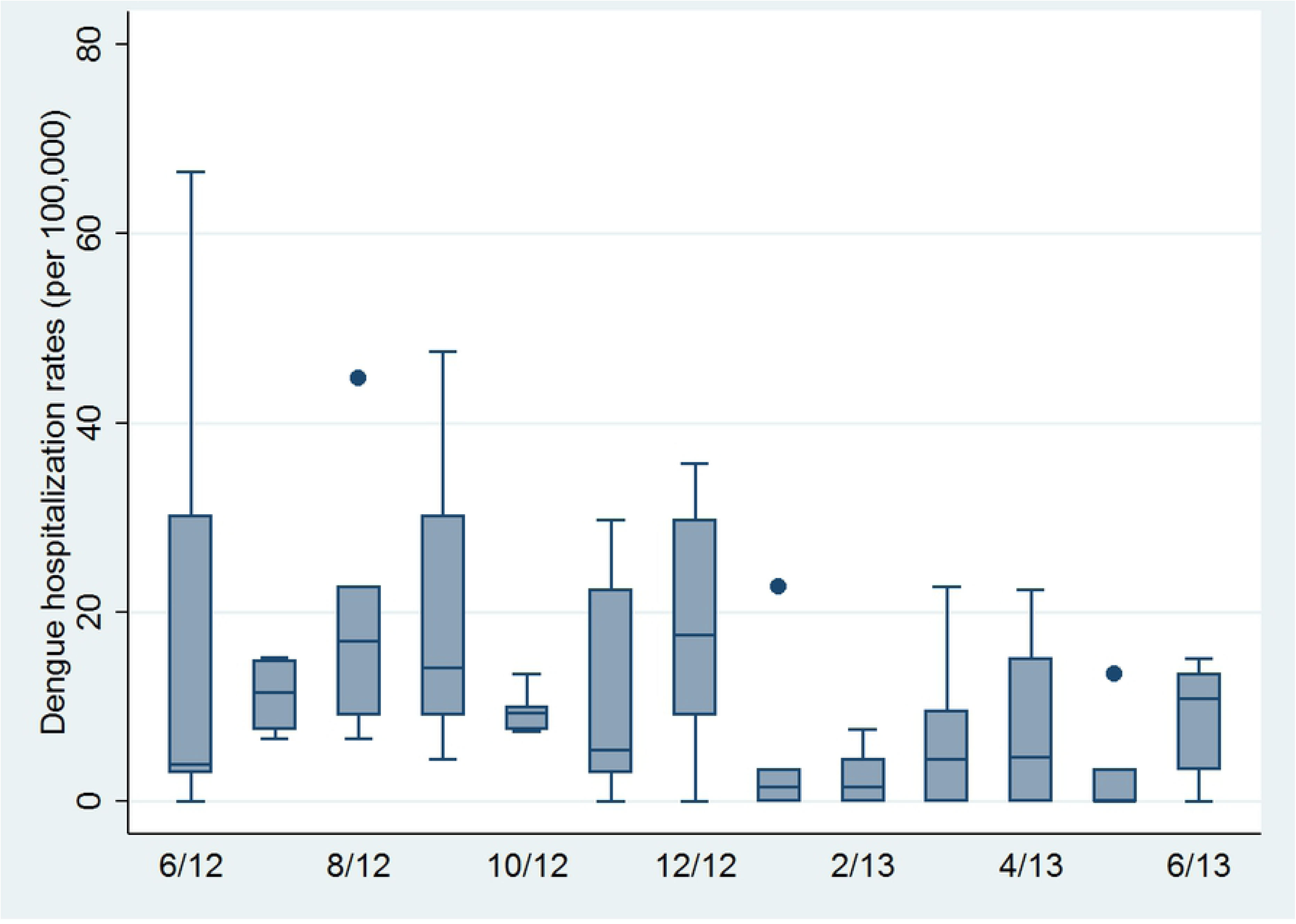
Monthly dengue hospitalization rates per 100,000 inhabitants for all wards, June 2012 – June 2013.

The dry season had a mean monthly hospitalization rate of 8.5 (95%CI 5.0, 12.1), whereas the rainy season had a mean of 12.8 (95%CI 8.6, 17.1) (p=0.13). Dengue rates were significantly associated with most of the meteorological variables at 0-, 1- and 2-month lags (Table 5a). Models with rainfall and relative humidity with a lag of 2-months had the lowest AIC. The only significant association between dengue hospitalization rates and vector indices was observed for PI with a 2-month lag (Table 5b).

### Mosquito avoidance behaviors by season

The percentage of people endorsing the use of mosquito avoidance behaviors was higher in the dry season (92.5%) than the rainy season (86.0%) (Table 6). The most commonly used method was the fan (80.0% in the dry season vs. 67.6% in the rainy season). Other common methods were using mosquito repellant coil and spray. The use of coils was significantly lower during the rainy season (47.2% in the dry season vs. 35.2% in the rainy season). Similarly, efforts to eliminate larvae were more frequent in the dry season than the rainy season (50.5% vs. 39.2%). The most common methods were cleaning and changing the water in the containers. The study participants engaged in this activity, as well as most of the other larval elimination methods, more frequently in the dry season than the rainy season.

**Table 6.**
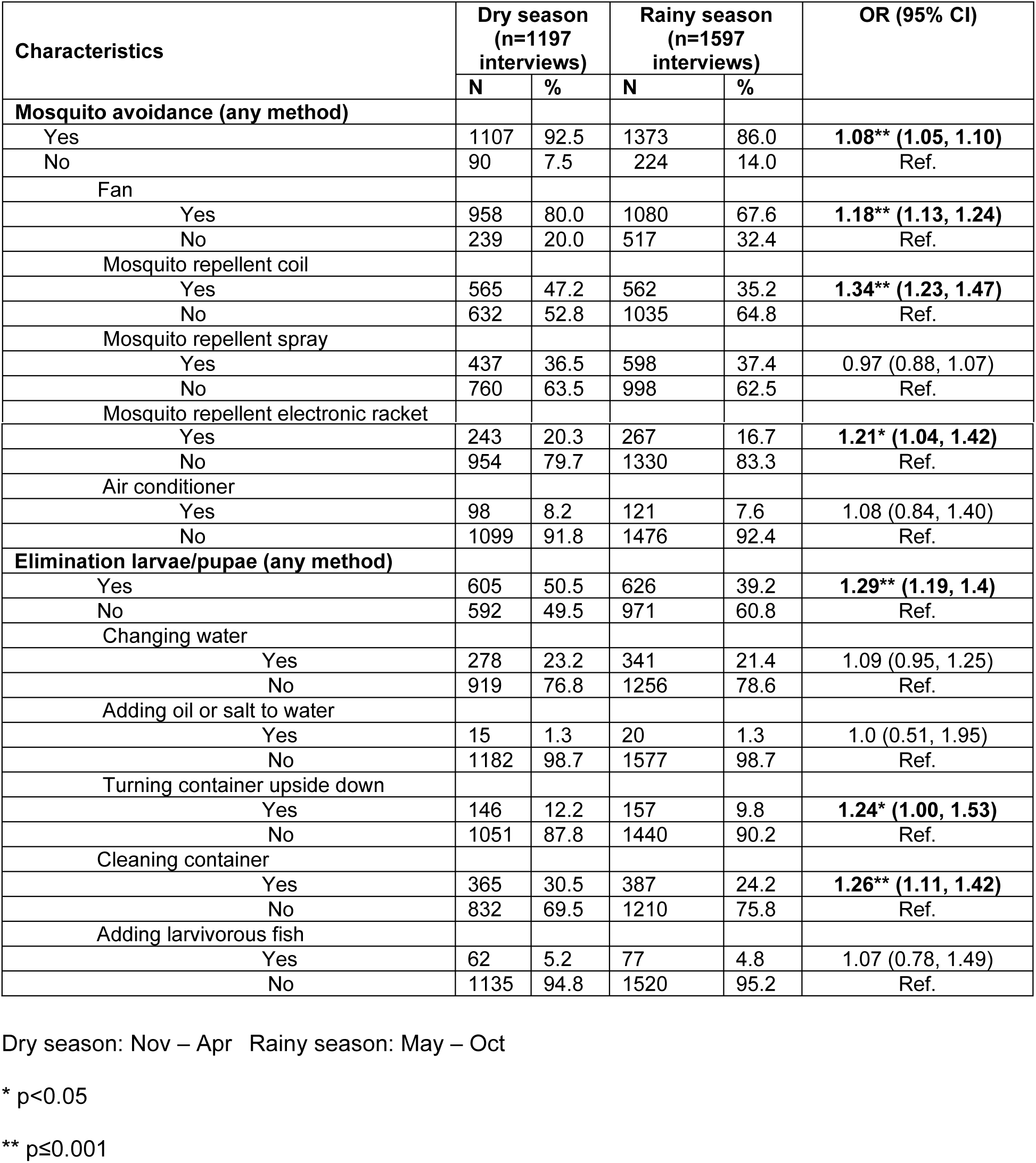
Mosquito avoidance behaviors and breeding site elimination strategies by season (McNemar test for paired proportions).

## Discussion

In this study, we describe associations between meteorological factors and a) vector indices, b) dengue hospitalization rates, and c) mosquito avoidance behaviors in a dengue endemic region of Vietnam. We show that relative humidity, in particular, is significantly associated with all of the vector indices, as well as the dengue hospitalization rate. The link between vector indices and dengue hospitalization rates is more tenuous, and only evident for the pupal index. Interestingly, the rainy season was associated with a reduction in a variety of behaviors that serve to reduce mosquito exposure or breeding, such as the use of fans, repellant coils and maintenance of water storage containers. Furthermore, we found that these water storage containers were the most important sources of *Ae. aegypti* pupa.

It has previously been shown that *Ae. aegypti* survival and fecundity are increased, and larval development accelerated, during periods of high humidity[14, 15]. Experimental studies have further demonstrated significantly higher dengue virus titers and an enhanced ability of the virus to proliferate within *Ae. aegypti* with increases in relative humidity[15]. We postulate that temperature in the Mekong Delta is largely within optimum range for vector breeding and viral dissemination throughout the year. A time-series analysis on regional differences in dengue incidence in Vietnam demonstrated that in Ho Chi Minh City (near Can Tho) where annual average temperature is 28°C, dengue incidence was positively associated with relative humidity, but negatively associated with temperature [16]. This stood in contrast to Hanoi (annual average temperature of 23.6°C), where the opposite was observed. This relationship between dengue transmission, temperature and humidity was modeled in a high-resolution profile of these factors across space in Peru [17]. Our findings are consistent with this study, which demonstrated that dengue transmission potential was dependent on the duration within an optimal temperature range and was amplified exponentially by high humidity [17].

In our study region, human behavior may also be contributing to the relationship between humidity, vector indices and dengue rates. Cleaning the large water storage containers may be quite cumbersome, especially when filled with water during the rainy season. However, it is exactly during the rainy season, when relative humidity and vector activity are high, that such maintenance is most needed. Pupal abundance is highest in these containers, accounting for the vast majority of *Ae. aegypti* pupae collected. Many households use these containers due to unreliable water supply, and they have been implicated previously as major sources of vector breeding[18]. Targeting productive containers for biological control with the copepod *Mesocyclops spp*. has been promoted in Northern and Central Vietnam[18] [19]. It has also been implemented in Southern Vietnam where disease burden is greatest, with great initial success[20]. However, subsequent studies noted challenges with the sustainability of this intervention in Southern Vietnam, due to the reluctance tointroduce organisms into drinking water [21]. In our study, we also found that larvivorous organisms such as copepods were rarely used.

Our multivariate model of larval density also highlights the importance of waste management and disheveled clothes in the house. The former has been borne out in other studies as well, which show that a lack of effective waste management increases the total number of *Ae. aegypti* breeding sites in disposable receptacles filled by rainfall [22–25]. In our study, however, disposable receptacles did not contribute much to pupal productivity, and the link between waste management and dengue infection remains undefined. The association between larval density and disorganized clothing in the house may be arising from the tendency of *Ae. aegypti* adult females to rest on clothing or on furniture below 1.5m while digesting a blood meal, preferably in bedrooms[26–28]. In fact, observing that *Ae. aegypti* females rest on clothing in dark locations while vacuum aspirating adult mosquitoes, Edman et al. fashioned resting boxes to mimic these conditions[29]. They found that boxes covered in black cloth were able to attract 20-70% of the adult *Ae. aegypti* population in a given house.

This study has several limitations. The entomological component of this study spans one year, which does not allow for the examination of longer term trends. Nonetheless, our findings with regards to the importance of relative humidity are consistent with time-series analyses for the region[7, 16]. Our use of dengue hospitalization rates no doubt represents only a small portion of dengue infections. While measuring dengue incidence would more accurately capture the outcome of interest, this endeavor requires active surveillance and is resource-intensive. Similarly, block-level indices of larval forms of *Ae. aegypti* do not tend to correlate as well with dengue transmission as the pupal and adult stage mosquito indicators[30]. Indeed, in our study the pupal index (with a 1-month lag period), but not the larval indices, were significantly associated with dengue hospitalizations.

There is emerging evidence that exposure to infected mosquitoes occurs not only in and around the household, but also in public spaces[31, 32]. Here we focused on measuring domiciliary vector indices exclusively, which does not fully capture dengue transmission risk for household members on a local scale. This may not have ultimately impacted our findings much, however, since we aggregated vector and dengue hospitalization data on the ward level, and further accounted for ward level heterogeneity in the mixed effects models.

Despite there still being many unanswered questions pertaining to the linkages between climate forecasts and projected changes in dengue transmission risk, there has been a shift in affected countries towards focusing on mitigation and adaptation. A recent special report issued by the Intergovernmental Panel on Climate Change indicated that low-lying coastal countries such as Vietnam are particularly vulnerable to impacts resulting from global warming of 1.5°C above pre-industrial levels[33]. Sea level rise and tidal flooding, rising temperatures, and extreme rainfall in southern Vietnam are already occurring[34], and downstream effects such as water shortages and salinity intrusion may already be impacting vector ecology. Current mitigation priorities in Vietnam include reducing carbon emissions, reforestation, improved water resources and waste management[35]. Some strategies to address these priorities may provide multiple health benefits, such as improving water supply and infrastructure, such that water storage will no longer be necessary. Prospectively measuring the impacts of such mitigation efforts on vector-borne disease indicators will provide valuable insight into the full extent of benefits conferred.

In conclusion, our study sought to link dengue and weather by examining multiple levels within the chain of causation, namely vector indices, pupal productivity, dengue hospitalization rates and mosquito avoidance and elimination measures. Our results indicate that relative humidity is a key weather variable in this area where temperatures are consistently within an optimal range for dengue transmission. We also found that large water storage containers are the source of the majority of *Ae. aegypti* pupae, and that these containers are maintained less frequently during the rainy season. Climate change projections forecast rising temperatures and flooding in this region of Vietnam. This will likely render this region vulnerable to water shortages, leading to more reliance on storing water near the domicile. Further studies are warranted on how these factors will influence not only dengue but also other the transmission risk of other arboviruses vectored by *Ae. aegypti.*

## Acknowledgements

We would like express our gratitude to the participants in this study. We are also indebted to Dr. Karoun Bagamian for producing the maps in this manuscript. We would like to express our appreciation for the Institute for Social and Environmental Transition (ISET), the Climate Change Coordination Office (CCCO), the Institute for Public Health in Ho Chi Minh, the Can Tho Department of Health, the Can Tho Preventive Medicine Center, Can Tho University of Medicine and Pharmacy, the Research Institute of Climate Change, Can Tho University, its collaborators in the health sector, and all of the individuals who have supported the implementation of this project. This project was funded by the Rockefeller Foundation.

## Supporting Information Legends

S1. Monthly dengue hospitalization rates in study districts in Can Tho, Vietnam, 2012-2013.

S2. Monthly larval indices and weather variables in Can Tho, Vietnam, 2012-2013.

S3. Illustration of water storage containers

